# A WD40 repeat-like protein pathway connects F-BOX STRESS INDUCED (FBS) proteins to the NIGT1.1 transcriptional repressor in Arabidopsis

**DOI:** 10.1101/2020.12.22.424016

**Authors:** Edgar Sepulveda-Garcia, Elena C Fulton, Emily V Parlan, Ashley A Brauning, Lily E O’Connor, Anneke A Fleming, Amy J Replogle, Mario Rocha-Sosa, Joshua M Gendron, Bryan Thines

**Affiliations:** Instituto de Biotecnología, Universidad del Papaloapan, Tuxtepec 68301, Mexico; Departamento de Biología Molecular de Plantas, Instituto de Biotecnología, Universidad Nacional Autónoma de México, Cuernavaca, Mor, 62250, Mexico; Biology Department, University of Puget Sound, Tacoma, WA 98416, USA; Department of Molecular, Cellular, and Developmental Biology, Yale University, New Haven, Connecticut 06511, USA

**Keywords:** F-box protein, SCF complex, stress response, transcription regulation, WD40 repeat-like protein

## Abstract

SCF-type E3 ubiquitin ligases use F-box (FBX) proteins as interchangeable substrate adaptors to recruit protein targets for ubiquitylation. FBX proteins almost universally have structure with two domains. A conserved N-terminal F-box domain interacts with a SKP protein and connects the FBX protein to the core SCF complex, while a C-terminal domain interacts with the protein target and facilitates recruitment. The F-BOX STRESS INDUCED (FBS) subfamily of four plant FBX proteins has atypical domain structure, however, with a centrally located F-box domain and additional conserved regions at both the N- and C-termini. FBS proteins have been linked to environmental stress networks, but no ubiquitylation target(s) or exact biological function has been established for this subfamily. We have identified two WD40 repeat-like proteins in Arabidopsis that are highly conserved in plants and interact with FBS proteins, which we have named FBS INTERACTING PROTEINs (FBIPs). FBIPs interact exclusively with the N-terminus of FBS proteins, and this interaction occurs in the nucleus. FBS1 destabilizes FBIP1, consistent with FBIPs being ubiquitylation targets of SCF^FBS^ complexes. Furthermore, we found that FBIP1 interacts with NIGT1.1, a GARP-type transcriptional repressor that regulates nitrate and phosphate starvation signaling and responses. Collectively, these interactions between FBS, FBIP, and NIGT1.1 proteins delineate a previously unrecognized SCF-connected transcription regulation module that works in the context of phosphate and nitrate starvation, and possibly other environmental stresses. Importantly, this work also identified two uncharacterized WD40 repeat-like proteins as new tools with which to probe how an atypical SCF complex, SCF^FBS^, functions via FBX protein N-terminal interaction events.

## INTRODUCTION

Essential plant processes, ranging from growth and development to stress responses, are controlled at the molecular level through selective protein degradation by the ubiquitin 26S proteasome system (UPS). Protein targets destined for removal are ubiquitylation substrates for E3 ubiquitin ligases, where one prevalent E3 ligase subtype is the SKP1-Cullin-F-box (SCF) complex (Hua and Vierstra, 2011). SCF complexes use an interchangeable F-box (FBX) protein subunit as a substrate adaptor to specifically interact with unique protein targets (Gagne et al., 2002; Sheard et al., 2010; Calderon Villalobos et al., 2012). FBX proteins almost universally have structure with two domains: an N-terminal F-box domain facilitates interaction with a SKP protein and the core SCF complex and a C-terminal domain interacts specifically with the target(s) (Gagne et al., 2002). This two-domain structure directly bridges core UPS components to precise protein targets under specific situations, and it places FBX proteins at a dynamic interface that regulates diverse cellular output pathways.

A very small number of FBX proteins across eukaryotes, however, deviate from this typical two-domain protein structure. Many of these atypical FBX proteins have a centrally located F-box domain, a C-terminal target interaction domain, and an additional protein interaction domain at the N-terminus (Jin, 2004; Wang et al., 2014; Lee et al., 2018). In humans, N-terminal domains can control subcellular localization (Matsumoto et al., 2011), bind to an accessory protein that assists with C-terminal targeting events (Spruck et al., 2001), or mediate regulatory interactions with other proteins (Jin, 2004; Kirk et al., 2008; Nelson et al., 2013). The only plant FBX proteins with established N-terminal interaction dynamics belong to the ZEITLUPE (ZTL), FLAVIN-BINDING KELCH REPEAT F-BOX1 (FKF1), and LOV KELCH PROTEIN2 (LKP2) subfamily, which regulates the circadian clock and flowering time (Yasuhara, 2004; Kim et al., 2007; Sawa et al., 2007; Zoltowski and Imaizumi, 2014; Lee et al., 2018). In addition to a central F-box domain, the ZTL/FKF1/LKP2 subfamily has an N-terminal blue-light sensing LOV domain and C-terminal kelch repeats (Zoltowski and Imaizumi, 2014), which are both used to recruit distinct ubiquitylation substrates (Más et al., 2003; Yasuhara, 2004; Song et al., 2014; Lee et al., 2018). The N-terminal LOV domain has additional roles that regulate FBX function through interaction with GIGANTEA (GI), which controls subcellular localization and protein stability (Kim et al., 2007; Sawa et al., 2007). Thus, across kingdoms, atypical FBX proteins with an N-terminal protein interaction domain, in addition to a C-terminal targeting domain, achieve expanded function by having further regulatory capacity and/or coordinating multiple cellular outputs through a dual targeting structure.

F-BOX STRESS INDUCED (FBS) proteins are a far less understood subfamily of four plant FBX proteins with atypical structure (Maldonado-Calderon et al., 2012; Sepulveda-Garcia and Rocha-Sosa, 2012; Gonzalez et al., 2017). FBS1 is the founding member of this FBX subfamily and is recognized for its broad biotic and abiotic stress responsive gene induction profiles (Maldonado-Calderon et al., 2012; Gonzalez et al., 2017). In FBS1, a centrally located F-box domain is flanked by two conserved regions present at the N- and C-termini, which do not match any known protein interaction domains or motifs (Maldonado-Calderon et al., 2012). FBS1 interacts with Arabidopsis SKP1 (ASK1) and can autoubiquitylate (Maldonado-Calderon et al., 2012; Sepulveda-Garcia and Rocha-Sosa, 2012), suggesting that it forms a functional SCF-type E3 ligase in vivo. At least five of thirteen Arabidopsis 14-3-3 regulatory proteins bind to FBS1 (Sepulveda-Garcia and Rocha-Sosa, 2012). However, because this interaction requires both the N-terminal region and the F-box domain of FBS1 (Sepulveda-Garcia and Rocha-Sosa, 2012), and ubiquitylation presumably requires an unhindered F-box domain to interact with the SKP subunit of the SCF complex (Hua and Vierstra, 2011), 14-3-3s are unlikely ubiquitylation targets. Furthermore, an inducible *FBS1* gene construct had no discernable effect on FBS1 interactor 14-3-3λ protein abundance (Sepulveda-Garcia and Rocha-Sosa, 2012). Importantly though, all five FBS1-interacting 14-3-3 proteins are negative regulators in Arabidopsis responses to cold and/or salt stress (Catala et al., 2014; van Kleeff et al., 2014; Zhou et al., 2014), which demonstrates another noteworthy cellular link between FBS1 and environmental stress response networks beyond the broad stress-inducible transcriptional regulation of *FBS1*.

More complete understanding of FBS protein function in plants has been stymied by two primary limitations. First, not knowing selective targeting relationship(s) between SCF^FBS^ complexes and their substrates has left FBS action on cellular output pathways completely enigmatic. Second, functional redundancy within this family has likely thwarted past efforts seeking to establish a biological function based on phenotype of Arabidopsis *fbs1* plants (Maldonado-Calderon et al., 2012; Gonzalez et al., 2017), but no evidence for redundancy exists to confirm this as an experimental barrier. Here, we identify two highly conserved WD40 repeat-like proteins that interact with multiple FBS family members in Arabidopsis, which we have named FBS INTERACTING PROTEINs (FBIPs). Interactions between all four FBS subfamily members and FBIP proteins occur in the nucleus, and interactions occur exclusively via the N-terminal domain of FBS proteins. FBIP1 also interacts in the nucleus with NIGT1.1, a DNA-binding GARP transcriptional repressor and key regulator of plant nitrate and phosphate signaling and starvation responses (Kiba et al., 2018; Maeda et al., 2018; Ueda et al., 2020a, 2020b). This FBS-FBIP-NIGT1.1 network of newly identified protein interactions strongly suggests the possibility that FBS family proteins use N-terminal interaction events to regulate stress genes and, in particular, genes involved in nitrate and phosphate starvation responses and signaling.

## METHODS

### Bioinformatics

Gene and protein sequences were obtained from The Arabidopsis Information Resource (www.arabidopsis.org). Protein sequences were aligned using T-COFFEE (www.ebi.ac.uk/Tools/msa/tcoffee) accessed through the European Bioinformatics Institute (EBI) website (www.ebi.ac.uk). WD40 repeat-like sequences were identified in FBIP1 and FBIP2 using the WD40-repeat protein Structures Predictor data base version 2.0 (WDSPdb 2.0; www.wdspdb.com) (Ma et al., 2019). Basic Local Alignment Search Tool (BLAST) and Position-Specific Iterative (PSI)-BLAST were accessed through the National Center for Biotechnology Information (NCBI) website (www.ncbi.nlm.nih.gov) and used to search the RefSeq database. Candidate protein interactors were identified by searching the SUBA4 database (www.suba.live) (Hooper et al., 2017).

### Gateway cloning

Gene-specific primers (Supplementary Table S1) were used with PCR to amplify coding sequences from pooled *Arabidopsis thaliana* (accession Col-0) cDNA. Amplicons were inserted into pENTR/D-TOPO vector (ThermoFisher Scientific) according to the manufacturer’s protocols. Genes were then transferred with LR Clonase II enzyme mix (ThermoFisher Scientific) into pCL112 or pCL113 (Zhu et al., 2008a) destination vectors for BiFC experiments, and into pGBKT7-GW (Addgene plasmid #61703) or pGADT7-GW (Addgene plasmid #61702) destination vectors for yeast two-hybrid experiments. Alternatively (Figure 3B), *FBS1* and *FBIP1* sequences were cloned into pBI770/pBI771 and tested for interaction, as done previously (Sepulveda-Garcia and Rocha-Sosa, 2012). Primers used to create *FBS1* truncation constructs are indicated in Supplementary Table S1.

### Yeast two-hybrid assays

*Saccharomyces cerevisiae* cells were grown, transformed, mated, and selected for by standard yeast protocols. Bait constructs (GAL4 DNA-binding domain, DBD) were transformed into Y2H Gold and prey constructs (GAL4 activation domain, AD) into Y187 strains by LiAc method (Takara Bio USA). Haploid strains were mated to produce diploid strains to test for interactions. Diploid strains were grown for 24 hours at 30 °C in liquid synthetic defined (SD) medium minus Trp/Leu (-TL) medium with shaking. Cells were then washed in sterile water, cell concentrations were adjusted to OD_600_ = 10^0^, 10^-1^, 10^-2^, 10^×3^, and 10 µL was spotted on SD -TL (control), SD minus Trp/Leu/His (-TLH), and SD minus Trp/Leu/His (-TLHA) selective plates. Plates were incubated for two days at 30 °C and then scanned to produce images.

### Bimolecular fluorescence complementation (BiFC)

Recombinant plasmids were transformed into *Agrobacterium tumefaciens* strain GV3101 (pMP90) by electroporation and selected under appropriate antibiotics. *A. tumefaciens* seed cultures were grown in LB with appropriate antibiotic selection for two days with shaking at 30 °C and then used to inoculate 50 mL LB containing appropriate antibiotics plus 10 µM acetosyringone and grown for an additional 24 hours. Cells were pelleted and resuspended in infiltration medium (10 mM MES, 10 mM MgCl_2_, 100 µM acetosyringone) and incubated for five hours with rocking at room temperature. Cells were pelleted a second time, resuspended in infiltration medium and appropriate nYFP/cYFP, H2B-RFP constructs were combined at a final OD_600_ of 1.0 for each test/control construct with suppressor strains (p19, γβ, PtoHA, HcPro) at a final OD_600_ of 0.5. *Nicotiana benthamia* leaves from four week-old plants were infiltrated by syringe with the *A. tumefaciens* mixes. The underside of whole leaf mounts was visualized using laser-scanning confocal microscopy three days after infiltration with a Nikon D-Eclipse C1 Confocal laser scanning microscope (Nikon Instruments) with either: 1) excitation at 488 nm with an emission band pass filter of 515/30, or 2) excitation at 561 nm with an emission band pass filter of 650 LP.

### Co-infiltration

*FBS1, FBIP1*, and *14-3-3λ* were cloned into pGWB17 (4X myc tag), pGWB14 (3X HA tag), or pGWB12 (VSVG tag) vectors (Nakagawa et al., 2007), respectively, using a Gateway strategy as above. Recombinant plasmids were transformed by electroporation into *A. tumefaciens* strain C58C1Rif/pGV2260. *A. tumefaciens* was grown to stationary phase in LB medium containing appropriate antibiotics plus 50 µg/ml acetosyringone. Bacteria were pelleted and washed with 10 mM MgCl_2_, and then resuspended in 10 mM MgCl_2_ and 150 µg/ml acetosyringone. Cell densities were adjusted to OD_600_ of 0.5. After 3 h of incubation, *A. tumefaciens* strains containing each construct were adjusted to varying concentrations and mixed with the same volume of an *A. tumefaciens* strain containing the viral suppressor p19, treated in the same way, but adjusted to OD_600_ of 1.0. The abaxial side of leaves from 3-4 week-old *N. benthamiana* were infiltrated with this bacterial suspension. After 3 days, leaf material was collected and immediately frozen in liquid N_2_ for protein extraction.

### Protein extraction and Western blotting

Approximately 100 µg of frozen tissue was homogenized in 200 µl of 1X Laemmli loading buffer plus 4 M urea, boiled 5 minutes and centrifuged at 10,000 × *g* for five minutes. 10 µl of the supernatant were loaded onto 8%, 10%, or 15% polyacrylamide gels and subjected to SDS-PAGE using standard protocols. Separated proteins were blotted onto a Hybond-P+ membrane (Amersham Pharmacia Biotech) using standard protocols, and then membranes were probed with anti-c-Myc, anti-HA antibody, or anti-VSVG antibodies (all from Sigma). Blots were developed using an alkaline phosphatase kit (BCIP/NBT kit; Invitrogen).

### AGI numbers

FBS1 (At1g61340), FBS2 (At4g21510), FBS3 (At4g05010), FBS4 (At4g35930), FBIP1 (At3g54190), FBIP2 (At2g38630), NIGT1.1 (At1g25550)

## RESULTS

### FBS protein interaction with ASK1

FBS1 is the founding protein of a four-member FBX protein subfamily (FBS1 – FBS4). FBS2 – FBS4, like FBS1, share non-canonical structure with a centrally located F-box domain and conserved regions at their N- and C-termini (Figure 1A). The conserved region at FBS N-termini spans approximately 20 residues, while the conserved region at the C-terminus encompasses about 35 (Figure 1A). FBS1 interacts with ASK1 and autoubiquitylates, indicating FBS1 likely participates in functional SCF complexes (Maldonado-Calderon et al., 2012; Sepulveda-Garcia and Rocha-Sosa, 2012). However, the ability of other FBS family members to interact with ASK proteins remains unknown, as does the possibility of functional redundancy among family members. To interrogate this possibility, all four FBS family members were tested as bait constructs (DBD, GAL4 DNA-binding domain) for interaction with ASK1 as prey (AD, GAL4 activation domain) under less stringent (-TLH) and more stringent (-TLHA) nutritional selection. Interactions were apparent between all four FBS family members on -TLH, although only very minimal growth was observed for FBS2 (Figure 1B). Only interactions between FBS1 and FBS4 with ASK1 were apparent under most stringent selection (-TLHA) (Figure 1B). Since Arabidopsis has 21 ASK proteins, it is possible the FBS proteins showing minimal partnering with ASK1 instead interact more strongly with other untested ASKs (Kuroda et al., 2012). These interactions show, however, that all FBS2 – FBS4 are viable candidates for functional SCF complex substrate adapters, like FBS1.

**Figure 1.**
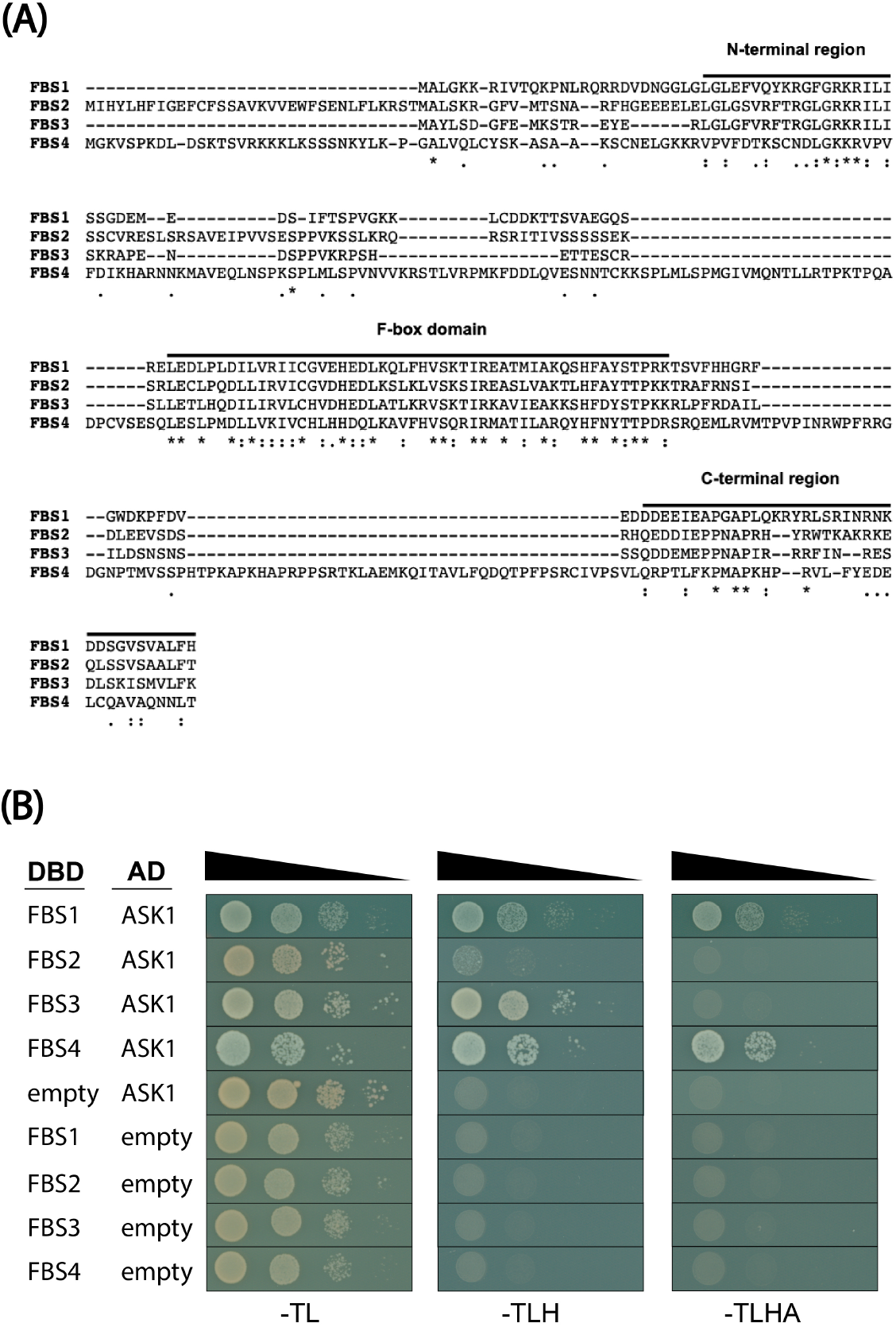
The F-BOX STRESS INDUCED (FBS) protein family. **(A)** Full-length protein sequence alignment of the four Arabidopsis FBS family members (FBS1 – FBS4) created with T-COFFEE sequence alignment program. Asterisks are fully conserved residues, colons are strongly conserved residue properties, and periods are weakly conserved residue properties. **(B)** FBS family interactions with ASK1 in yeast two-hybrid assays. Diploid yeast strains with indicated test constructs as bait (DBD) and prey (AD) were grown in liquid culture, diluted (OD_600_ = 10^0^, 10^×1^, 10^×2^, 10^×3^), and spotted on SD medium minus Trp/Leu (-TL), minus Trp/Leu/His (-TLH), and minus Trp/Leu/His/Ade (-TLHA).

### Identification of a new FBS1 interactor

In addition to ASK1, the only known FBS1 interacting proteins are 14-3-3 proteins (Sepulveda-Garcia and Rocha-Sosa, 2012). However, because interaction dynamics are not consistent with ubiquitylation of 14-3-3 proteins by SCF^FBS1^ (Sepulveda-Garcia and Rocha-Sosa, 2012), we sought additional FBS1 interactors as candidate targets that could connect FBS proteins to biological processes. Two additional related proteins were identified as partners for FBS1, which we have named FBS INTERACTING PROTEINs (FBIPs). FBIP1 (At3g54190) was identified in the same yeast two-hybrid screen that found 14-3-3 proteins as FBS1 interactors (Sepulveda-Garcia and Rocha-Sosa, 2012). FBIP1 is also listed as an FBS1 interactor by the SUBA4 database (www.suba.live) from previous high-throughput protein-protein interaction (PPI) screening (Arabidopsis Interactome Mapping Consortium et al., 2011; Hooper et al., 2017). FBIP1 is 467 residues in length and is a member of the transducin / WD40 repeat-like superfamily of proteins. WD40 repeats typically form a β-propeller domain that acts as a scaffold in mediating protein-protein or protein-DNA interactions (Jain and Pandey, 2018). Seven putative WD40 repeat-like sequences were predicted in FBIP1 by the WD40-repeat protein Structures Predictor database version 2.0 (WDSPdb 2.0) (Ma et al., 2019), although these predictions fall into the low confidence category (Figure 2). A second FBIP protein (At2g38630) was identified in the Arabidopsis genome by BLAST search, which we have named FBIP2. Protein sequence identity and similarity between FBIP1 and FBIP2 are just over 91% and 96%, respectively (Figure 2).

**Figure 2.**
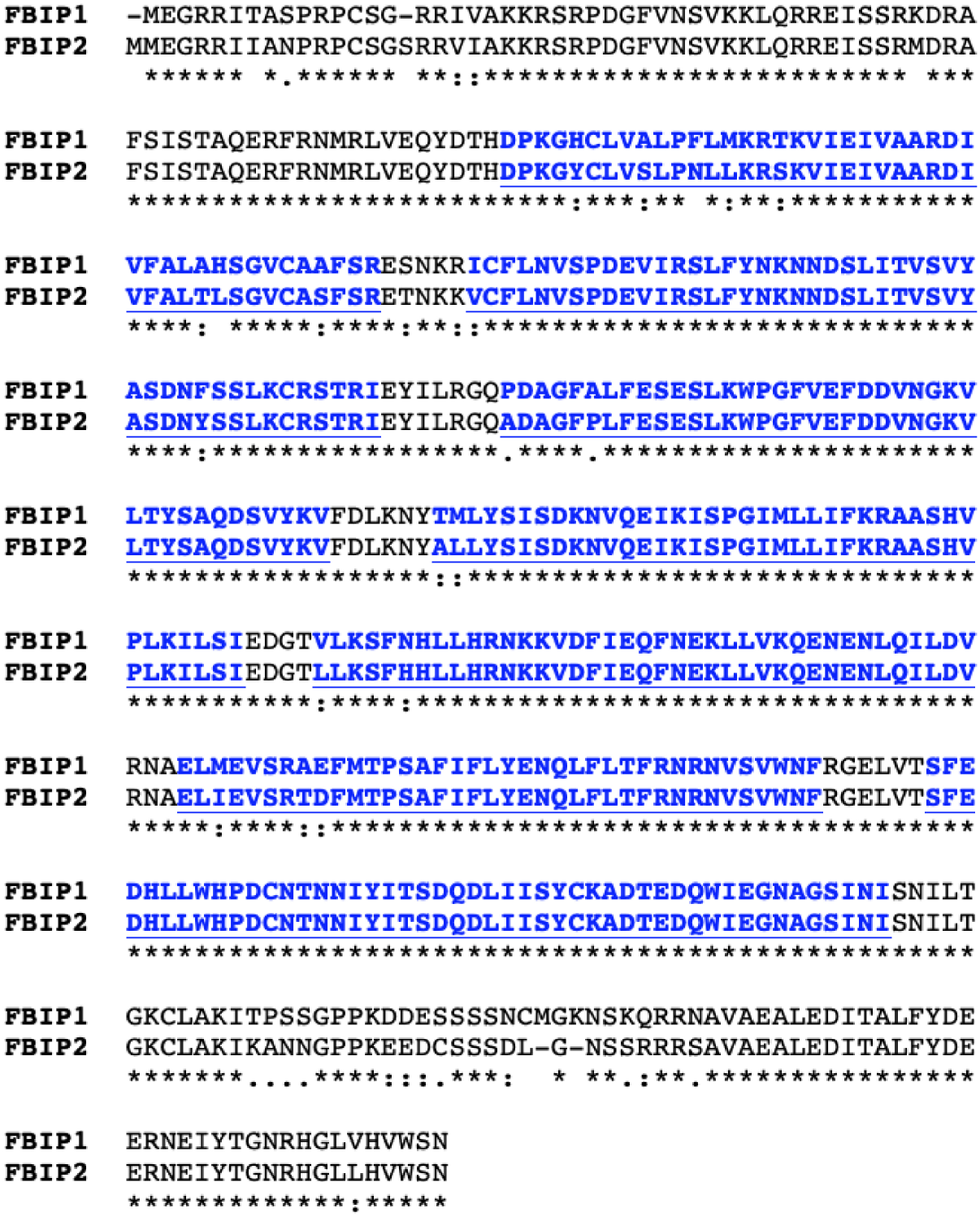
FBS INTERACTING PROTEIN (FBIP) sequence features. Full-length protein sequence alignment of the two Arabidopsis FBIP family members created with T-COFFEE sequence alignment program. Blue indicates locations of seven WD40-like repeat sequences predicted by the WD40-repeat protein Structure Predictor version 2.0 (WDSPdb 2.0). Asterisks are fully conserved residues, colons are strongly conserved residue properties, and periods are weakly conserved residue properties.

We gained no additional insight about FBIP function using various bioinformatics resources. Other than putative WD repeat-like sequences, no sequence features were identified using various domain or motif prediction programs. BLAST and PSI-BLAST searches with FBIP1 and FBIP2 sequences failed to identify additional significant hits in Arabidopsis. We did, however, find very highly conserved FBIP sequences throughout the plant kingdom, including in bryophytes (the top BLAST hit in *Physcomitrella patens* is about 77% identical and 85% similar to *Arabidopsis* FBIP1). By investigating AtGenExpress ATH1 array data sets (Schmid et al., 2005; Kilian et al., 2007; Goda et al., 2008), we found that *FBIP1* is constitutively expressed in most tissues and organs of Arabidopsis, and throughout its life cycle, but we found no conditions where *FBIP1* is more highly expressed compared to other conditions. *FBIP2* is not represented on the ATH1 array.

### FBS interactions with FBIPs

We confirmed that full-length FBS1 and FBIP1 interact by yeast two-hybrid analysis. Interaction between FBS1 and FBIP1 elicited growth in yeast strains on both less stringent (-TLH) and more stringent (-TLHA) nutritional selection, and FBS1 yielded growth with FBIP2 on -TLH (Figure 3A). Family-wide interactions between each FBS protein and the two FBIP proteins were also assessed (Figure S1). Growth was also observed for FBS3 and FBIP1, but not with FBS2 or FBS4. No additional interactions were observed with FBIP2. Collectively, yeast two-hybrid results suggest that FBS1 and FBIP1 might be the primary FBS-FBIP protein interaction pair, or possibly bind with strongest affinity, but that some other family-wide interactions might be possible.

**Figure 3.**
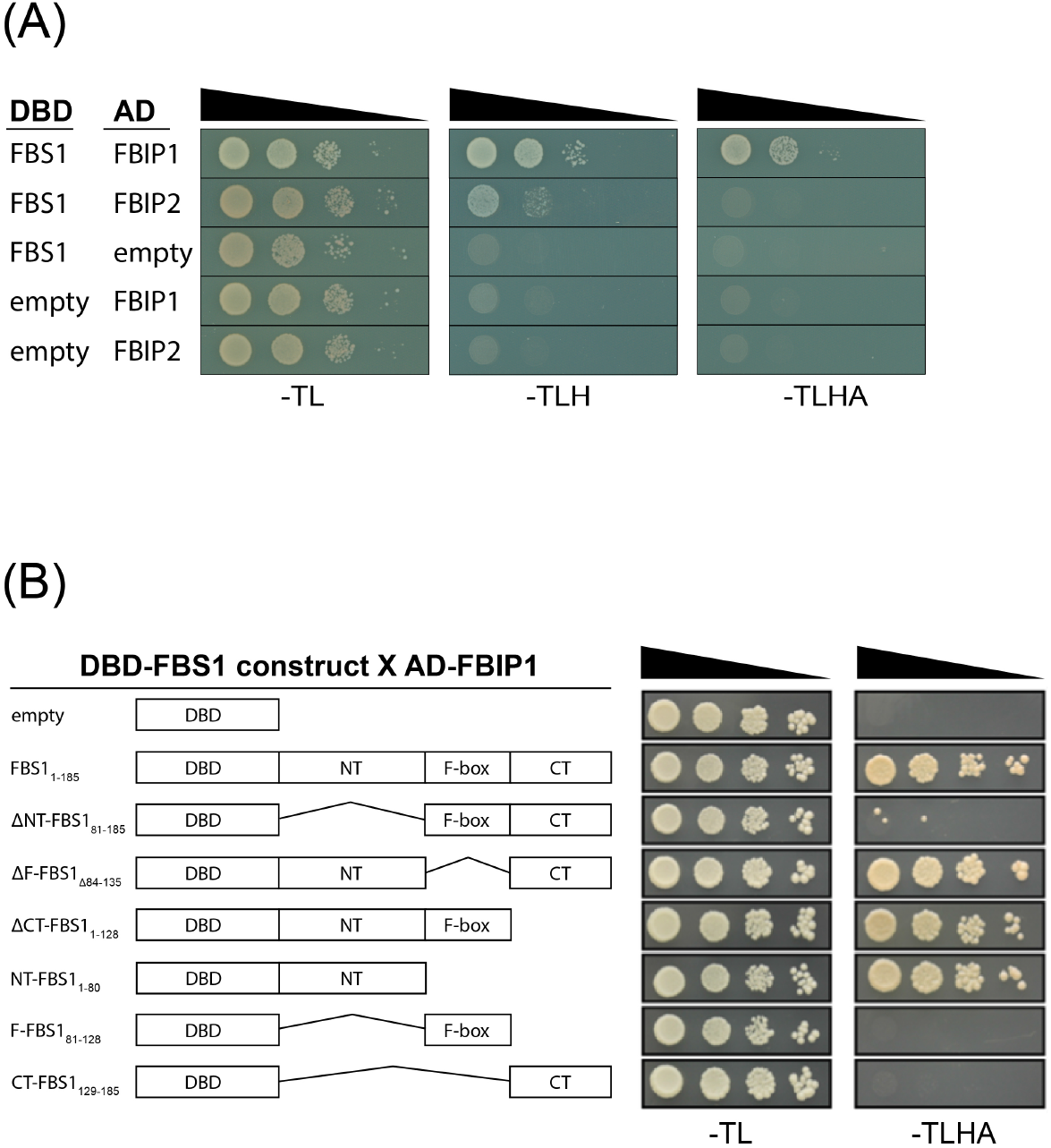
Yeast two-hybrid (Y2H) interactions between FBS1 and FBIP proteins. **(A)** Full-length FBS1 interactions with full-length FBIP1 and FBIP2. Diploid yeast strains with indicated test constructs as bait (DBD) and prey (AD) were grown in liquid culture, diluted (OD_600_ = 10^0^, 10^×1^, 10^×2^, 10^×3^), and spotted on SD medium minus Trp/Leu (-TL), minus Trp/Leu/His (-TLH), and minus Trp/Leu/His/Ade (-TLHA). **(B)** Truncated FBS1 bait (DBD) construct interaction with full length FBIP1 prey (AD). Amino acid deletions are indicated on left.

FBS proteins have two regions of unknown function outside of the F-box domain and, presumably, at least one of these interacts with a target. In order to determine which parts of FBS1 are important for FBIP1 interaction, we created truncated versions of FBS1 with the N-terminal (NT), F-box, or C-terminal (CT) regions removed in different combinations and tested under stringent (-TLHA) selection (Figure 3B). Removing the N-terminal region (ΔNT-FBS1_81-185_) abolished the ability of FBS1 to interact with FBIP1, while removal of the F-box domain (ΔF-FBS1_Δ84-135_) or C-terminal region (ΔCT-FBS1_1-128_) did not. The FBS1 N-terminal region (NT-FBS1_1-80_) in combination with full-length FBIP1 yielded growth on -TLHA, indicating that the FBS1 N-terminal domain alone is sufficient to mediate this interaction.

In the conserved N-terminal domains of FBS1 and FBS2 we found an overlapping LXLXL sequence (Figure 1A), which is the most prominent form of an EAR motif found in many different types of transcriptional regulators (Kagale and Rozwadowski, 2011; Shyu et al., 2012). The EAR motif mediates interaction with the WD40 repeat-containing protein TOPLESS (TPL) and TOPLESS RELATED (TPR) co-repressor proteins (Long, 2006; Pauwels et al., 2010; Causier et al., 2012). We considered whether this LXLXL sequence in the N-terminal region of FBS1 might: 1) function as a canonical EAR motif to interact with TOPLESS, and/or 2) if it could be important for mediating interactions with FBIPs. However, substituting all three leucine residues for alanine in FBS1 did not alter its interaction with FBIP1, and FBS1 did not interact with TPL (both as bait or as prey) in our yeast two-hybrid system.

### FBS interactions with FBIP occur in the nucleus

We next used bimolecular fluorescence complementation (BiFC) to test FBS interaction with FBIP in plants and determine where the interaction occurs in a cell. FBS and FBIP family proteins were expressed in *Nicotiana benthamiana* leaves as C-terminal fusions to either N-terminal (nYFP) or C-terminal (cYFP) halves of yellow fluorescent protein (YFP). In multiple independent experiments, YFP fluorescence was observed for pairings between FBS1 and FBIP1 and FBIP2 (Figure 4). This YFP signal co-localized with that of a co-infiltrated H2B-RFP construct, which localizes exclusively in the nucleus (Wang et al., 2013), and shows that interactions between FBS1 and FBIP proteins also occur in the nucleus. Similar experiments found that FBS2 – FBS4 also interact with FBIP1 in the nucleus (Supplementary Figure S2). We observed interactions for FBS3 and FBS4 with FBIP2 (Supplementary Figure S3), although we note that these interactions were more variable in number of YFP positive nuclei across independent replicates. We did not observe any interactions between FBS2 and FBIP2. All FBS and FBIP fusion protein constructs were tested as pairs with empty nYFP or cYFP vectors, and in all pairings we were unable to detect any fluorescent signal similar FBS/FBIP test pairs (Supplementary Figure S4). These findings show that in plants FBS proteins participate in family-wide interactions in the nucleus.

**Figure 4.**
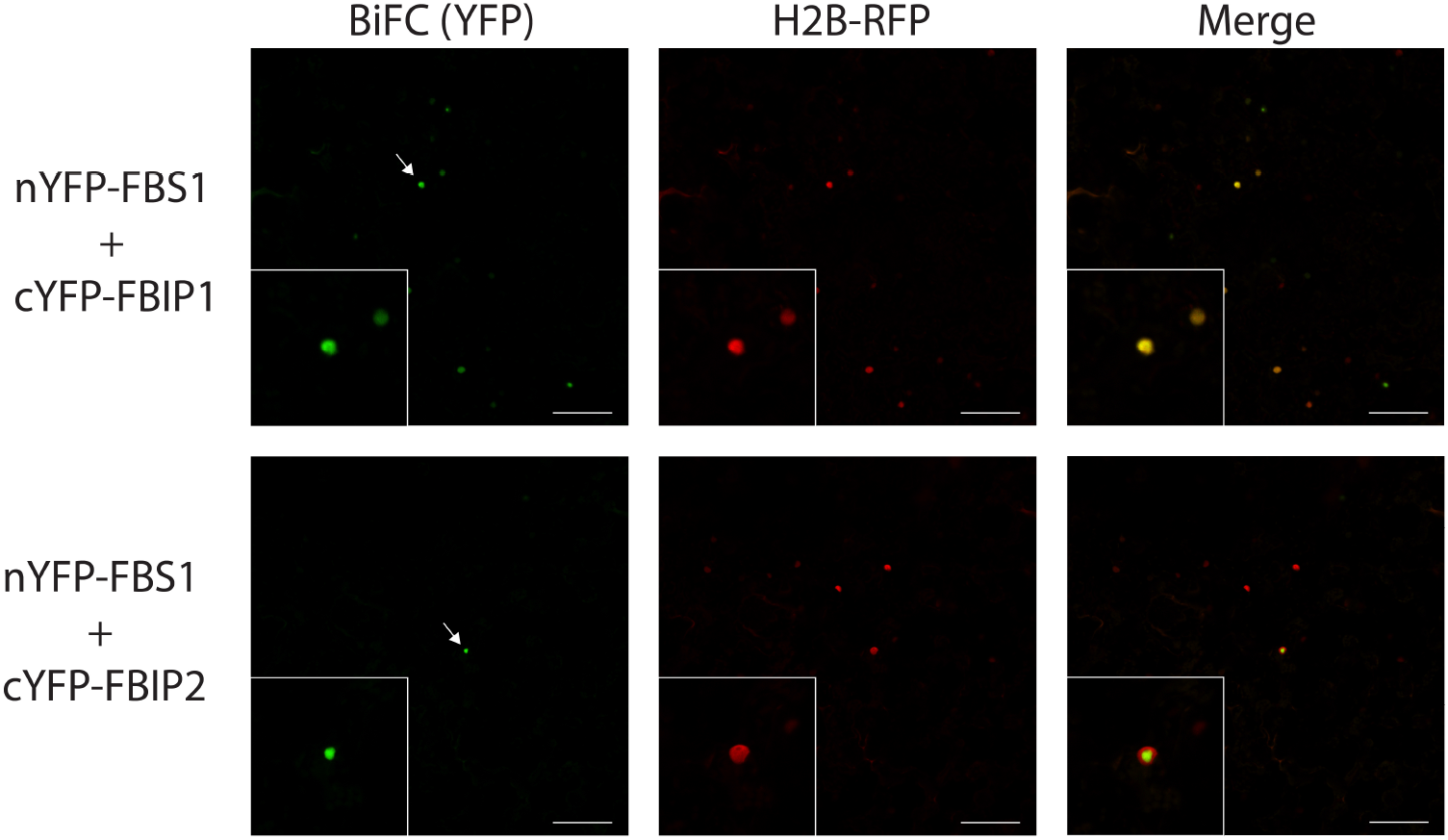
Bimolecular fluorescence complementation (BiFC) interactions between FBS1 and FBIP proteins. Laser-scanning confocal microscopy of *N. benthamiana* epidermal cells expressing N-terminal nYFP- or cYFP-tagged FBS1 and FBIP proteins. FBS1 interactions with FBIP1 (top row) or FBIP2 (bottom row) are visualized on BiFC yellow channel (YFP, left column). A co-expressed H2B-RFP (as nuclear marker) is visualized on red channel (RFP, middle column) and YFP/RFP images are overlaid (Merge, right column). Arrow indicates selected nuclei in expanded inset image. Scale bar = 100 µm

### FBS1 destabilizes FBIP1

With interaction established between multiple FBS and FBIP protein pairs, we next asked if the protein abundance relationship between FBS1 and FBIP1 is consistent with FBIP1 being a ubiquitylation target of SCF^FBS1^. If a protein is ubiquitylated by a particular SCF complex and subsequently degraded by the 26S proteasome, then increasing abundance of the F-box component typically increases in vivo targeting and decreases substrate abundance (dos Santos Maraschin et al., 2009). We therefore tested the effects of varying FBS1 protein levels on FBIP1 abundance in our *N. benthamiana* expression system by co-infiltrating *Agrobacterium* harboring these test constructs in different relative concentrations. Increasing the presence of FBS1 protein resulted in a corresponding decrease in FBIP1 protein abundance by Western blot analysis (Figure 5). In comparison, when FBS1 abundance was increased relative to co-infiltrated 14-3-3λ in an identical setup we did not observe any decrease in 14-3-3λ abundance as the amount of expressed FBS1 was increased (Supplementary Figure S5). This finding is congruous with previous observations that FBS1 and 14-3-3 interactions are not consistent with targeting (Sepulveda-Garcia and Rocha-Sosa, 2012). Therefore, because the abundance of FBIP1 decreases in an FBS1-dependent manner, we conclude that FBIPs are viable candidates for SCF^FBS1^ ubiquitylation targets.

**Figure 5.**
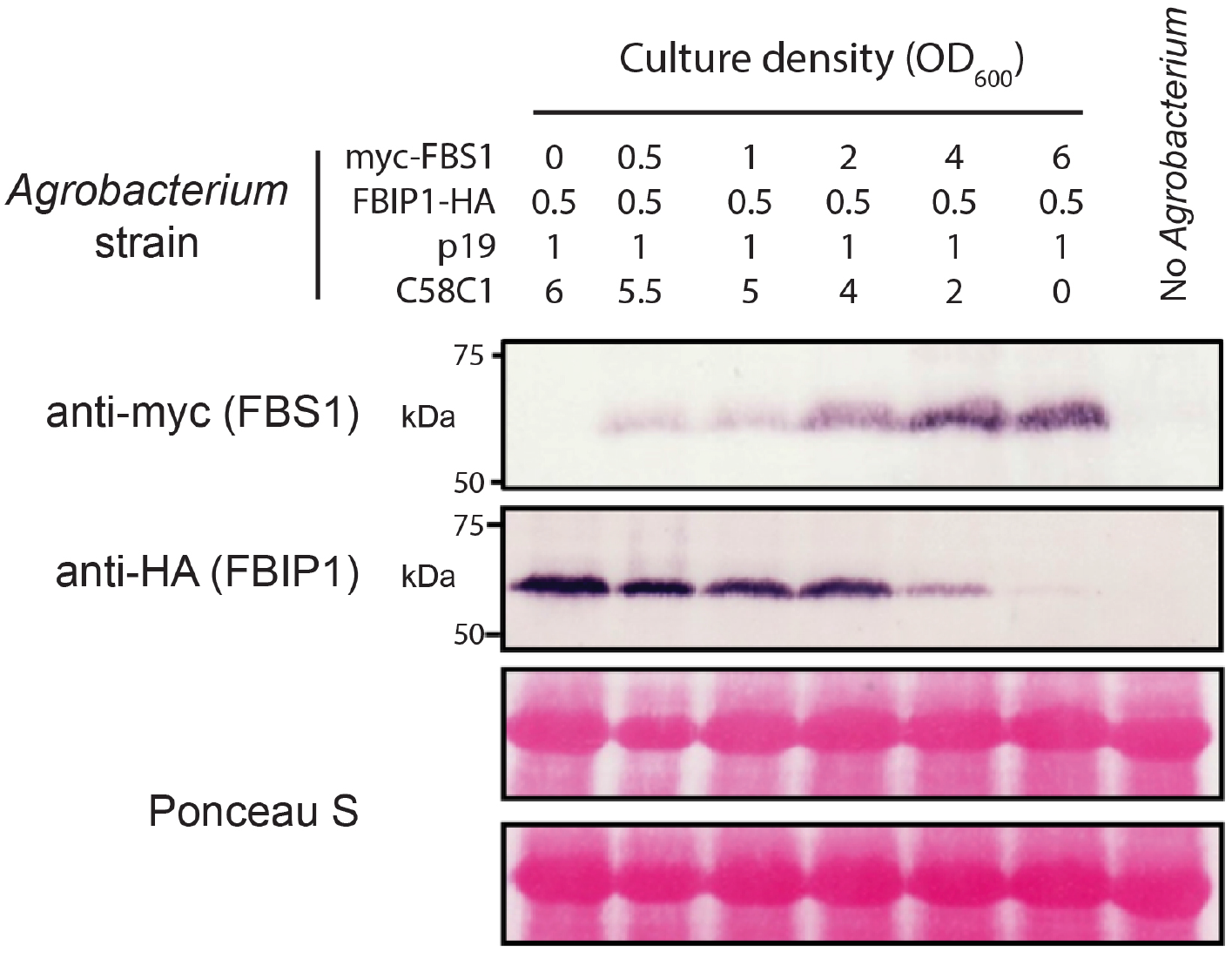
FBS1 influence on FBIP1 protein abundance in plants. *N. benthamiana* leaves were infiltrated with *Agrobacterium* (C58C1) strains to express tagged proteins. *Agrobacterium* mixes contained varying cell densities of strains harboring expression constructs (myc-FBS1 and/or FBIP1-HA), a suppressor protein (p19), or untransformed cells. Total protein was isolated from leaves three days after infiltration, separated by SDS-PAGE, transferred, and probed with antibodies against myc (top row, FBS1) or HA (second row, FBIP1). Bottom two rows show Ponceau S staining of the major subunit of Rubisco from the same two blots as a loading control.

### Interaction between FBIP1 and NIGT1.**1**

Interaction between FBS and FBIP protein families represents a newly recognized link between an SCF complex with stress inducible components (ie. *FBS1* gene expression; 14-3-3 interaction) and a potential targeting output. However, without knowing the precise biological function of FBIP proteins we cannot know the consequences of FBS-FBIP interactions, nor can we strongly connect FBS1 to an exact cellular pathway. Therefore, we examined protein interactions in the SUBA4 database for FBIP1, with particular consideration for our findings that FBS and FBIP interactions occur in the nucleus. One protein reported to interact with FBIP1 was Nitrate-Inducible GARP-type Transcriptional Repressor 1.1 (NIGT1.1/HHO3; At1g25550). NIGT1.1 is a DNA-binding transcriptional repressor and a central regulator of gene expression programs that coordinate nitrate (NO_3_^-^) and phosphate (PO_4_^3-^) signaling and starvation responses in plants (Kiba et al., 2018; Maeda et al., 2018; Ueda and Yanagisawa, 2019; Ueda et al., 2020a, 2020b). We tested this predicted interaction between FBIP1 and NIGT1.1 in yeast two-hybrid assays and observed growth on both less stringent (-TLH) and more stringent (-TLHA) conditions (Figure 6A). In BiFC, both FBIP1 and FBIP2 interacted with NIGT1.1 in the nucleus, as demonstrated by co-localization with H2B-RFP (Figure 6B). These interactions link FBS proteins through FBIP proteins to a DNA-binding transcriptional repressor, which suggests that at least one function of FBS proteins is to directly regulate gene expression programs that relate to environmental conditions (ie. nitrate and phosphate macronutrient availability).

**Figure 6.**
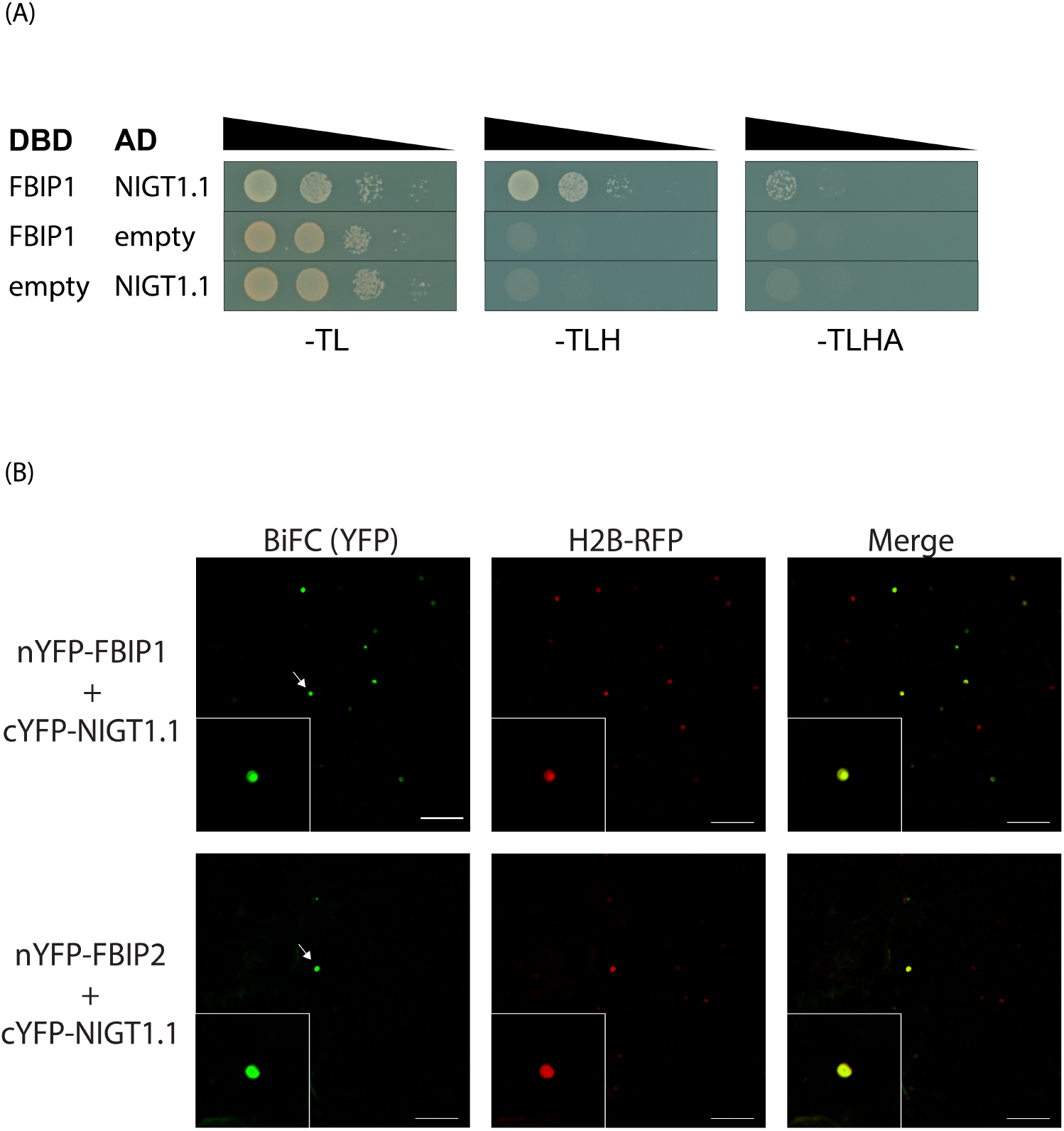
FBIP interactions with transcriptional repressor NIGT1.1. **(A)** Interaction between full-length FBIP1 and full-length NIGT1.1 in yeast two-hybrid assays. Diploid yeast strains with indicated test constructs as bait (DBD) and prey (AD) were grown in liquid culture, diluted (OD_600_ = 10^0^, 10^×1^, 10^×2^, 10^×3^), and spotted on SD medium minus Trp/Leu (-TL), minus Trp/Leu/His (-TLH), and minus Trp/Leu/His/Ade (-TLHA). **(B)** Laser-scanning confocal microscopy of *N. benthamiana* epidermal cells expressing N-terminal nYFP- or cYFP-tagged FBIP and NIGT1.1 proteins. NIGT1.1 interactions with FBIP1 (top row) or FBIP2 (bottom row) are visualized on BiFC yellow channel (YFP), left column). A co-expressed H2B-RFP (as nuclear marker) is visualized on red channel (RFP, middle column) and YFP/RFP images are overlaid (Merge, right column). Arrow indicates selected nuclei in expanded inset image. Scale bar = 100 µm

## DISCUSSION

Prior work with the FBS subfamily strongly alluded to its role in plant stress responses (Maldonado-Calderon et al., 2012; Sepulveda-Garcia and Rocha-Sosa, 2012; Gonzalez et al., 2017), but detailed understanding was limited by the unknown nature of ubiquitylation target(s) and by possible redundancy within the *FBS* gene family. Here, we have identified a pair of WD40 repeat-like superfamily proteins, FBIP1 and FBIP2, that both interact with FBS family proteins. Family-wide interactions between FBS and FBIP proteins in plants indicate that redundancy issues likely need to be circumvented before genetic approaches will yield full insight into *FBS* gene function based on phenotype analysis. Nonetheless, FBIP proteins are strong candidates for SCF^FBS^ ubiquitylation targeting. FBIP interaction with NIGT1.1, a key regulator of nitrate responsive genes, directly links FBS proteins to nuclear and transcription regulatory processes (Figure 7). Collectively, the FBS-FBIP-NIGT1.1 module is a new protein interaction network in which to understand regulation of stress genes by an SCF-type E3 ligase (Figure 7). Finally, FBIP and FBS interactions provide new context with which to investigate FBX protein N-terminal events, and to further understand how this unique subfamily of FBX proteins might couple N-terminal and C-terminal events to integrate cellular outputs to help plants maintain resilience under environmental stress.

**Figure 7.**
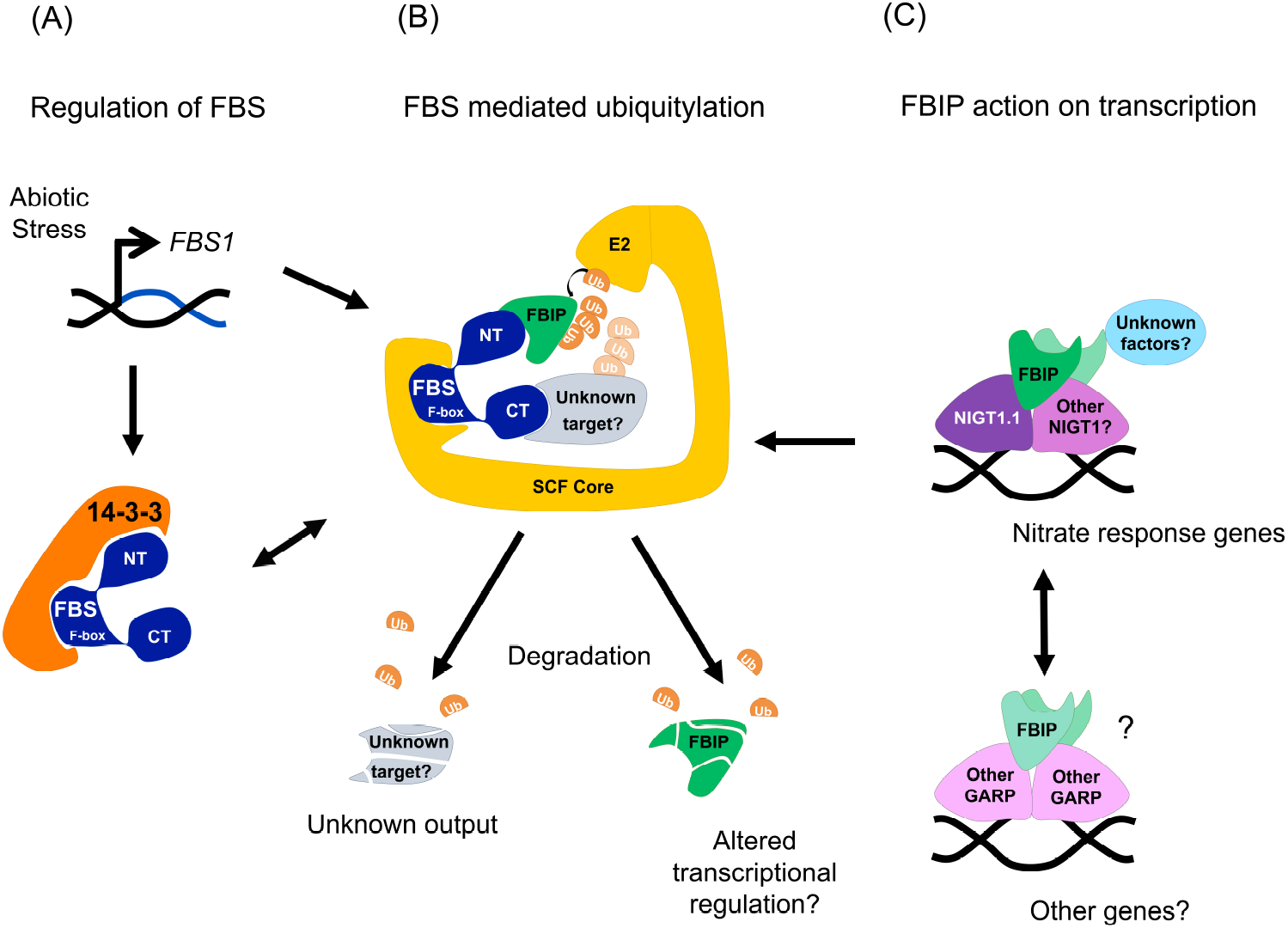
Integration of FBS proteins in a plant stress network. **(A)** Stress regulates FBS function through *FBS1* gene induction and possible control imposed by 14-3-3 proteins, which are negative regulators of abiotic stress responses. **(B)** SCF^FBS^ complexes ubiquitylate (Ub) FBIP through FBS N-terminal (NT) interactions and also target an unknown protein by FBS C-terminal (CT) interactions. Targets are degraded by the 26S proteasome leading to cellular changes under stress conditions. **(C)** NIGT1.1 dimerizes with other NIGT1 transcription factors and binds promoter regions of nitrate responsive genes. FBIP interacts NIGT1.1, and possibly with other NIGT1 and GARP-type transcription factors to influence their activity. Action by FBIP might influence in vivo dimerization, recruit additional gene regulation factors, alter DNA binding, or carry out some other function.

### The molecular function of FBIP proteins

Our findings point to a direct role for FBS proteins in gene regulation, but knowing this with certainty will require understanding the molecular function of FBIP proteins. Some plant nuclear-localized WD40 repeat proteins have direct actions in transcription regulation (Causier et al., 2012; Ke et al., 2015; Long and Schiefelbein, 2020) or chromatin modification (Li et al., 2007; Zhu et al., 2008b; Mehdi et al., 2016), and knowledge of these roles should inform hypotheses and future work. For example, TOPLESS (TPL) is a well-studied WD40 repeat-containing co-repressor protein that interacts with multiple transcriptional complexes acting in diverse pathways (ie. auxin, jasmonate, development) and recruits chromatin modifying enzymes to repress gene expression (Krogan et al., 2012; Wang et al., 2013). TRANSPARENT TESTA GLABRA 1 (TTG1), another WD40 repeat protein, serves as a scaffold and mediates different combinations of bHLH and R2R3-type MYB transcription factors to regulate flavonoid metabolism and various developmental processes (Lloyd et al., 2017; Long and Schiefelbein, 2020). Considering these established roles for WD40 repeat proteins in nuclear events, a few possibilities seem readily apparent for FBIPs in the context of NIGT1.1-mediated transcription regulation. First, FBIPs could recruit additional proteins that either enable or inhibit the transcriptional repression activity of NIGT1.1, potentially by interfacing with chromatin modifying enzymes, such as histone deacetylases (Wang et al., 2013). Second, as NIGT1.1 itself belongs to a subfamily of four NIGT1 transcription factors that dimerize (Yanagisawa, 2013; Ueda et al., 2020b), it is possible that FBIPs in some way mediate in vivo pairings and are functionally analogous to TTG1. Furthermore, as there are 56 GARP-type transcriptional repressors in Arabidopsis (Safi et al., 2017), it is possible that FBIP proteins could interact with some of these other regulators to exert broader effects on gene regulation beyond nitrate- and phosphate-dependent processes. We note that other GARP family transcription factors regulate ABA- and JA-responsive genes (Merelo et al., 2013), and so past work showing that *FBS1* impacts genes responsive to these two stress hormones is consistent with this notion (Gonzalez et al., 2017). Future efforts will be aimed at understanding the full spectrum of interactions between the two FBIP proteins and other GARP family transcription factors, with special focus on the NIGT1 subfamily, as well as whether FBIPs interact with additional proteins that may assist in gene regulation.

### FBIPs as candidate ubiquitylation targets

A number of important questions surround the consequence of FBIP proteins as FBS interactors, but hypotheses for immediate future work are equally apparent. Knowing that SCF complexes in some unique contexts ubiquitylate targets via FBX protein N-terminal interactions (Lee et al., 2018), and that FBS1 appears to destabilize FBIP1 (Figure 5), a leading hypothesis is that FBIP proteins are bona fide ubiquitylation substrates for SCF^FBS^. Rigorous assessment of in vivo interaction dynamics between SCF^FBS^ complexes and FBIP proteins, and whether interaction stimulates ubiquitylation-dependent degradation of FBIP proteins, will be critical lines of inquiry in future work. Given the constitutive gene expression profile of *FBIP1* across publicly accessible transcriptome data sets, it could be that FBIP proteins are components of a stress-response system that is triggered at the post-translational level. An obvious following question, then, is whether FBIP proteins are degraded in response to changing environmental conditions and, if so, whether some factor (ie. post-translational modification) stimulates SCF^FBS^ association with FBIP proteins under these conditions. The idea that additional in vivo factors or modification mediates FBS/FBIP interaction is consistent with notion that we observed more family-wide interactions in our in plant BiFC experiments compared to yeast two-hybrid.

With current understanding, however, we cannot completely exclude the possibility that FBIP proteins are not targets, but instead serve an alternative function that enables (or inhibits) FBS action. An idea with precedence is that FBIP proteins are accessories that recruit other proteins as ubiquitylation targets. For example, in Arabidopsis, KAI2 and D14 interact with FBX protein MAX2 in SCF^MAX2^ complex to mediate ubiquitylation of SMXL transcription factors (Wang et al., 2020). In humans, Cks1 directly associates with the N-terminus of FBX protein Skp2 to direct SCF^Skp2^ interaction with ubiquitylation target p27 in human cell cycle regulation (Spruck et al., 2001; Skaar et al., 2013). A parallel, but intimately connected, line of questioning involves identifying an FBS C-terminal region-interacting protein that we presume to exist. Knowing this additional putative interactor may aid in addressing important aspects of FBIP function, and future work can investigate the coordination of higher order complex assembly and/or possible situations of dual targeting and co-occurring processes.

### FBS proteins are new tools with which to probe regulation of nitrate/phosphate starvation responses

Nitrate and phosphate are two indispensable macronutrients, but their abundances are highly variable in most environments. The subfamily of four NIGT1 transcription factors directly regulates hundreds of nitrate responsive genes by: 1) helping to elicit a quick-pulse response to nitrate under some regulatory contexts (Ueda and Yanagisawa, 2019), or 2) control sustained diminished expression in other regulatory contexts (Medici et al., 2015; Ueda and Yanagisawa, 2019). Nitrate uptake and assimilation by plants is intimately coordinated with that of phosphate, and at least some regulatory events that accomplish this at the gene expression level occur through NIGT1 activities (Ueda and Yanagisawa, 2019). Though functional relationships between FBS, FBIP, and NIGT1.1 proteins are not yet known, recent work with NIGT1 proteins and their regulation nitrate and phosphate responsive gene networks gives invaluable experimental context for future work (Kiba et al., 2018; Maeda et al., 2018; Ueda et al., 2020a). Coupling Arabidopsis genetic resources related to *FBS* and *FBIP* genes to those of *NIGT1*.*1* will likely advance our understanding of how these factors work together, for example whether FBIP proteins have a positive effect on NIGT1.1 (and other NIGT1 family proteins), to accomplish regulation of nitrate-responsive transcriptional processes in various environmental contexts (ie. cold or salt stress). Furthermore, as both *NIGT1* and *FBS1* are very rapidly induced by their respective stress-inducing situations (Maldonado-Calderon et al., 2012; Sawaki et al., 2013; Gonzalez et al., 2017), understanding how these factors work together may help further define temporal priorities and resource management in nitrogen acquisition and other parts of stress responses. Taken together, harnessing *FBS* and *FBIP* genes will present new opportunities by which to understand how plants integrate and manage nitrate and phosphate stresses with other stress conditions.

Different stress response pathways do not work in isolation (Rasmussen et al., 2013), but are coordinated with one another to collectively contribute to comprehensive health of plants under duress. However, much remains to be learned about the integration of different pathways. Given its broad biotic and abiotic stress-triggered induction, as well as its stress hormone responsiveness (Maldonado-Calderon et al., 2012; Gonzalez et al., 2017), *FBS1* may act in a common cellular pathway or process that is more universally harnessed to aid compromised, or otherwise challenged, plant cells. Further support for this notion comes from the fact that FBS1 interacts with multiple 14-3-3 proteins that work at least in both salt and cold stresses (Sepulveda-Garcia and Rocha-Sosa, 2012; Catala et al., 2014; van Kleeff et al., 2014; Zhou et al., 2014). The mechanistic connection delineated by an FBS/FBIP/NIGT1 module may connect a more globally induced environmental stress response to a nitrate uptake/assimilation program mediated by NIGT1 and co-acting proteins. In fact, nitrogen, in particular the nitrate and ammonia forms, enhances plant performance in various forms of abiotic stress, as it is required for *de novo* synthesis of various metabolites and proteins with protective properties (Zhang et al., 2018; Rohilla and Yadav, 2019; Li et al., 2020). In seeming contrast, however, some abiotic stress-responsive transcriptional networks naturally limit expression of genes central to nitrogen uptake and assimilation (Goel and Singh, 2015). These observations underscore the notion that there is still much to learn about the complexities of these gene regulatory networks and physiological processes acting in broader stress contexts. This work, including the subsequent hypotheses it generates, provides a new mechanistic framework in which to assess how an atypical SCF complex may coordinate cellular stress pathways, including those acting in nitrate and phosphate uptake and assimilation, through transcription regulation events.

## AUTHOR CONTRIBUTIONS

ESG, ECF, EVP, AAB, LEO, AAF, AJR, and BT conducted the experiments. All authors designed the experiments, analyzed the data, and approved the final version of the manuscript. BT wrote the manuscript.

## ACKNOWLEDGEMENTS

We thank David Somers (The Ohio State University) for the H2B-RFP construct, Frank Harmon (University of California at Berkeley / USDA Plant Gene Expression Center) for the yeast two-hybrid vectors, Faride Unda (University of British Columbia) for the BiFC vectors, and Ruirui Huang and Vivian Irish (Yale University) for the yeast two-hybrid TOPLESS constructs. We also thank Marisa Weiss for her help with cloning *NIGT1*.*1* and Andreas Madlung (University of Puget Sound) for critical reading of the manuscript and other helpful discussions. Finally, we would like to thank Michal Morrison-Kerr (University of Puget Sound) for her indispensable help in supporting Puget Sound undergraduate research students.

