## Supplementary materials for "A WD40 repeat-like protein pathway connects F-BOX STRESS INDUCED (FBS) proteins to the NIGT1.1 transcriptional repressor in Arabidopsis"

### SUPPLEMENTAL MATERIALS

### SUPPLEMENTARY FIGURE LEGENDS

**Supplementary Figure S1. Yeast two-hybrid FBS1 – FBS4 interactions with FBIP1 and FBIP2.** FBS2 – FBS4 family interactions with FBIP1 and FBIP2 in yeast two-hybrid assays. Diploid yeast strains with indicated test constructs as bait (DBD) and prey (AD) grown in liquid culture, were diluted ($OD_{600}$ = $10^0$, $10^{-1}$, $10^{-2}$, $10^{-3}$), and spotted on SD medium minus Trp/Leu (-TL), minus Trp/Leu/His (-TLH), and minus Trp/Leu/His/Ade (-TLHA).

**Supplementary Figure S2. Bimolecular fluorescence complementation (BiFC) interactions between FBS1 – FBS4 and FBIP1.** Laser-scanning confocal microscopy of *N. benthamiana* epidermal cells expressing N-terminal nYFP- or cYFP-tagged FBS and FBIP1 proteins. FBS2 (top row), FBS3 (middle row), and FBS4 (bottom row) interactions with FBIP1 are visualized on BiFC yellow channel (YFP, left column). A co-expressed H2B-RFP (as nuclear marker) is visualized on red channel (RFP, middle column) and YFP/RFP images are overlaid (Merge, right column). Arrow indicates selected nuclei in expanded inset image. Scale bar = 100 μm

**Supplementary Figure S3. Bimolecular fluorescence complementation (BiFC) interactions between FBS1 – FBS4 and FBIP2.** Laser-scanning confocal microscopy of *N. benthamiana* epidermal cells expressing N-terminal nYFP- or cYFP-tagged FBS and FBIP1 proteins. FBS2 (top row), FBS3 (middle row), and FBS4 (bottom row) interactions with FBIP2 are visualized on BiFC yellow channel (YFP, left column). A co-expressed H2B-RFP (as nuclear marker) is visualized on red channel (RFP, middle column) and YFP/RFP images are overlaid (Merge, right column). Arrow indicates selected nuclei in expanded inset image. Scale bar = 100 μm

**Supplementary Figure S4. YFP channel positive and negative controls.** Laser-scanning confocal microscopy of *N. benthamiana* epidermal cells expressing GFP behind a CaMV 35S promoter (left), or nYFP- and cYFP fusion protein test constructs in combination with empty vectors. Note that image second from the right is that same as in Figure 4, for comparison, and the image on the far right is the same image as in Figure 4 but zoomed out. Scale bar in all images = 100 μm

**Supplementary Figure S5. FBS1 influence on 14-3-3 λ protein abundance in plants.** *N. benthamiana* leaves were infiltrated with *Agrobacterium* (C58C1) strains to express tagged proteins. *Agrobacterium* mixes contained varying cell densities of strains harboring expression constructs (myc-FBS1 and/or 14-3-3 λ-VSVG), a suppressor protein (p19), or untransformed cells. Total protein was isolated from leaves three days after infiltration, separated by SDS-PAGE, transferred, and probed with antibodies against myc (top row, FBS1) or VSVG (second row, 14-3-3 λ). Bottom two rows show Ponceau S staining of the major subunit of Rubisco from the same two blots as a loading control.

### SUPPLEMENTARY TABLES AND FIGURES

**Supplementary Table S1. Primer sequences used for cloning**

| **Function** | **Name** | **Sequence (5' → 3')** |
|---|---|---|
| Gateway cloning for yeast two-hybrid and BiFC | FBS1_5' | CACCATGGCATTGGGGAAGAAAAGAATCG |
| | FBS1_3' | TCAGTGGAATAGAGCCACTGAGAC |
| | FBS2_5' | CACCATGATCCATTATCTCCATTTCA |
| | FBS2_3' | TCATGTAAACAAAGCCGCAG |
| | FBS3_5' | CACCATGGCGTATTTGAGTGATG |
| | FBS3_3' | TCATTTAAACAATACCATAGAGATCTTCGACAAATC |
| | FBS4_5' | CACCATGGGGAAGGTATCTCCAAAG |
| | FBS4_3' | TCAGGTGAGGTTGTTTTGAGC |
| | FBIP1_5' | CACCATGGAAGGAAGAAGGATTAC |
| | FBIP1_3' | TCAATTGGACCACACATGAAC |
| | FBIP2_5' | CACCATGATGGAAGGAAGAAGAATCAT |
| | FBIP2_3' | TCAATTAGACCAGACATGGAG |
| Cloning for yeast two-hybrid (Figure 3B) | SalFBS1_5' | TGCAGG**GTCGAC**ATATGGCATTGGG |
| | NotFBS1_3' | GTATCGATA**GCGGCCGC**TCAGTGGAATA |
| | SalFBS1 _5' | TGCAGG**GTCGAC**ATATGGCATTGGG |
| | NotNtFBS1_3' | A**GCGGCCGC**TCACTCTCGACTCTGACC |
| | SalFFBS1_5' | C**GTCGAC**TTGAAGATCTTCCCCTAGATA |
| | NotFFBS1_3' | CC**GCGGCCGC**TCAGGTGTACTATACGC |
| | SalCtFBS1_5' | C**GTCGAC**GTACACCTCGGAAAACCTCGG |

**Supplementary Figure S1**

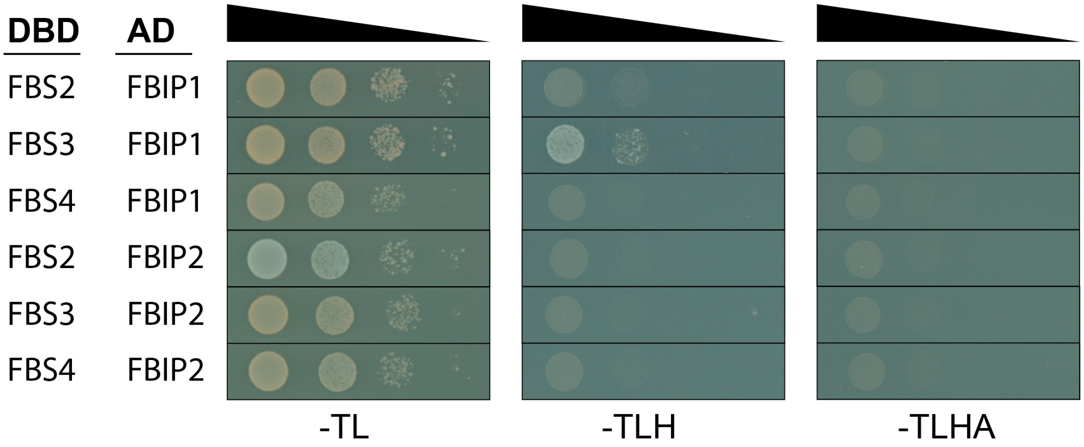

**Supplementary Figure S2**

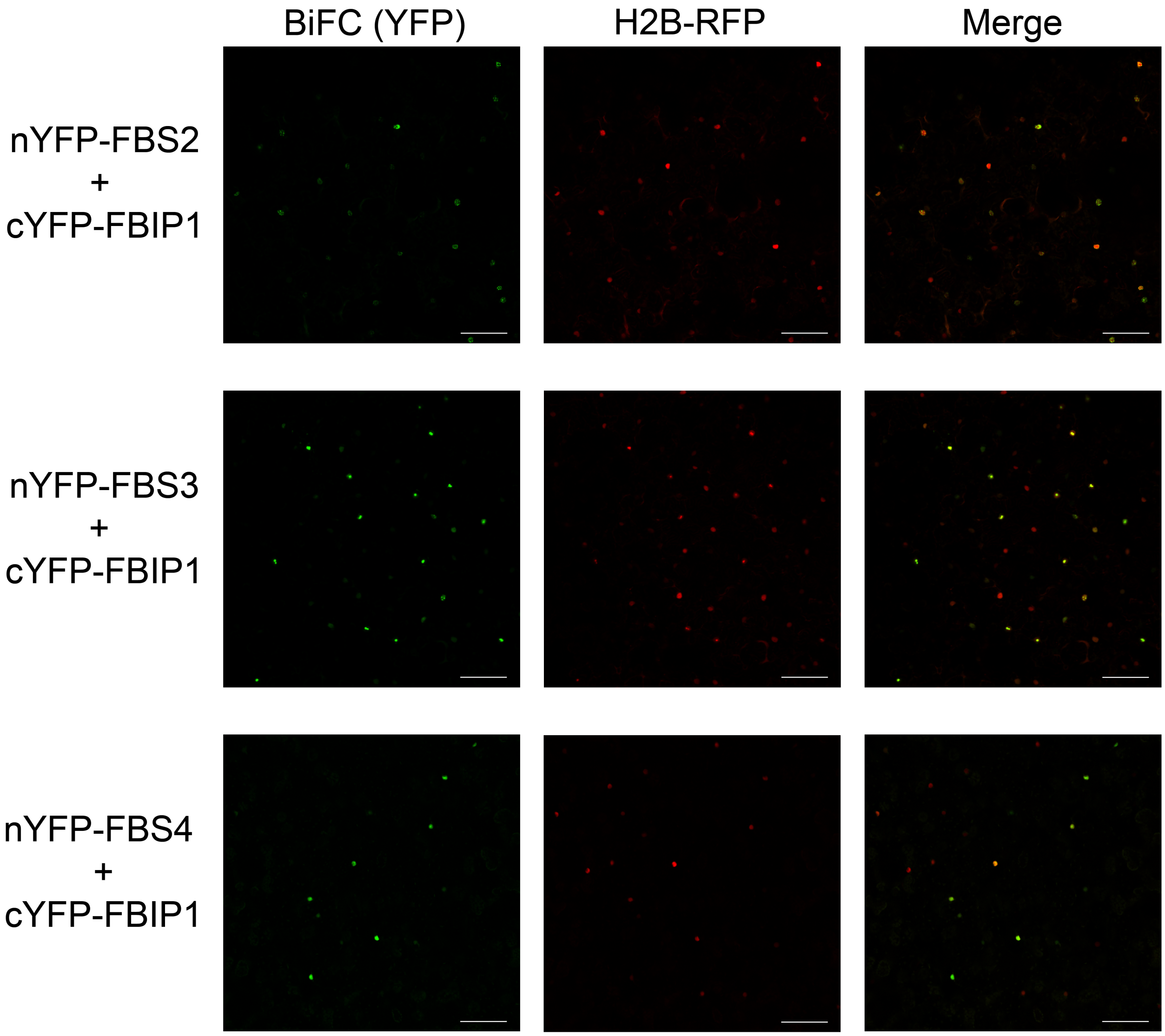

**Supplementary Figure S3**

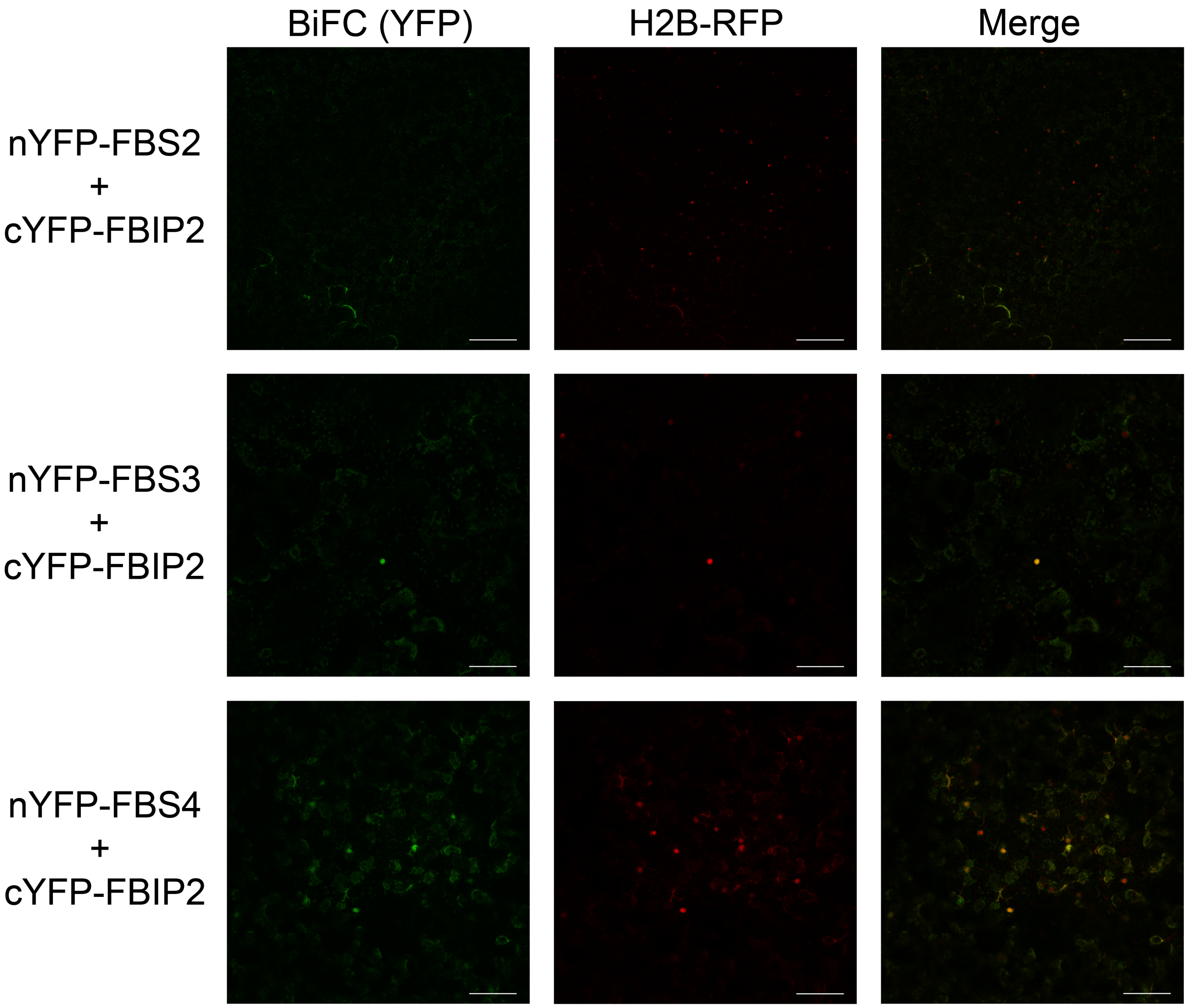

**Supplementary Figure S4**

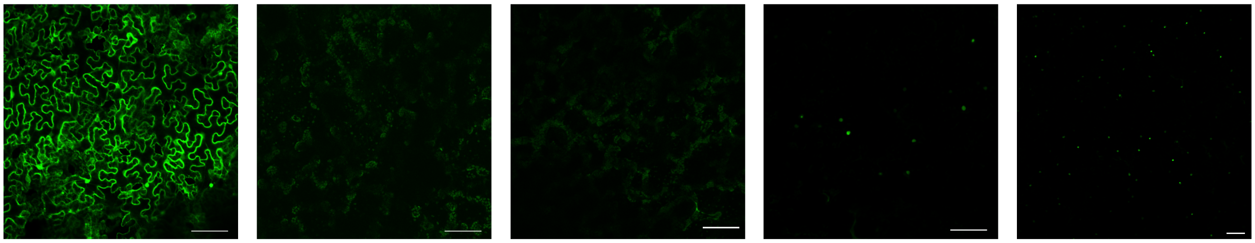

GFP | nYFP-empty + cYFP-FBIP1 | nYFP-FBS1 + cYFP-empty | nYFP-FBS1 + cYFP-FBIP1 | nYFP-FBS1 + cYFP-FBIP1

160 **Supplementary Figure S5**

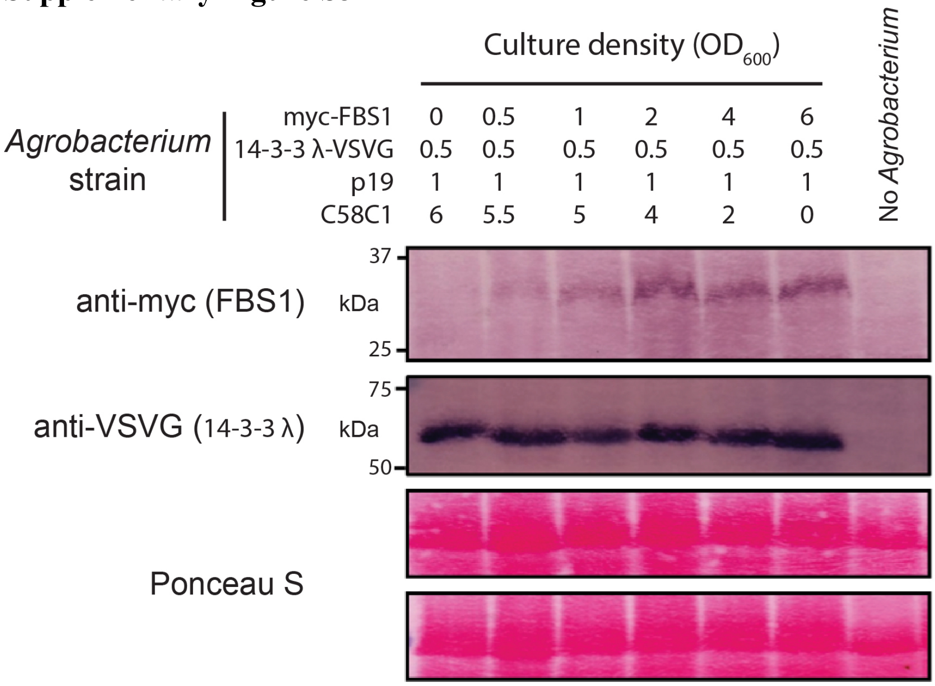


161
